# Keratinocyte desmoglein 1 regulates the epidermal microenvironment and tanning response

**DOI:** 10.1101/423269

**Authors:** Christopher R. Arnette, Jennifer L. Koetsier, Joshua A. Broussard, Pedram Gerami, Jodi L. Johnson, Kathleen J. Green

**Affiliations:** Department of Pathology, Northwestern University, Feinberg School of Medicine, Chicago, Illinois, 60611; Department of Dermatology, Northwestern University, Feinberg School of Medicine, Chicago, Illinois, 60611; Department of Pediatrics, and the Northwestern University, Feinberg School of Medicine, Chicago, Illinois, 60611; Lurie Comprehensive Cancer Center, Northwestern University, Feinberg School of Medicine, Chicago, Illinois, 60611

**Keywords:** Desmosomal cadherin, Ultraviolet, Melanocyte, Cytokine

## Abstract

Coordinated responses to environmental stimuli within the keratinocyte:melanocyte niche are poorly understood. Desmoglein 1 (Dsg1), a keratinocyte-specific desmosomal cell-cell adhesion protein with emerging signaling roles, is reduced by ultraviolet light radiation. Loss-of-function Dsg1 mutations elevate keratinocyte cytokines in Severe dermatitis, multiple Allergies, and Metabolic wasting (SAM) syndrome. We asked whether Dsg1 regulates keratinocyte:melanocyte paracrine communication to induce the tanning response. Dsg1-silenced keratinocytes increased *Pro-opiomelanocortin* mRNA and cytokine secretion. Melanocytes treated with conditioned media from Dsg1-silenced keratinocytes exhibited increased *Mitf* and *Trp1* mRNA, melanin secretion, and dendrite length. Inhibiting the melanocyte pigment-associated melanocortin 1 receptor reduced pigment secretion in response to Dsg1-deficient conditioned media. Melanocytes incorporated into Dsg1-deficient human skin equivalents relocalized suprabasally, reminiscent of early melanoma pagetoid behavior. Dsg1 decreased in keratinocytes surrounding dysplastic nevi and early melanoma, but not benign nevi. We posit Dsg1 controls keratinocyte:melanocyte communication through paracrine signaling, which goes awry upon Dsg1 loss in melanoma development.

## INTRODUCTION

The epidermis is a multi-layered organ composed of several cell types that form a barrier against environmental, pathogenic, chemical, and physical assaults and against water loss. Keratinocytes (KCs), the major cell type of the epidermis, undergo a program of differentiation whereby they transit out of the proliferative basal cell layer and exit the cell cycle, giving rise to suprabasal layers (spinous, granular, and stratum corneum) (Jensen and Proksch, 2009; Proksch *et al.*, 2008). Melanocytes (MCs), the pigment producing cells of the epidermis, reside within the basal layer of the skin. Through the extension of dendrites, MCs interact with KCs in a ratio of roughly 36 KCs: 1 MC, forming the KC:MC pigmentary unit (Nguyen and Fisher, 2018; Weiner *et al.*, 2014). KCs and MCs directly associate through cadherin-based adhesive structures and indirectly communicate via paracrine signaling (Lee and Herlyn, 2007; Mescher *et al.*, 2017; Serre *et al.*, 2018). The release of secreted factors (cytokines, chemokines, and growth factors) from KCs and other resident skin cells results in modulation of MC proliferation, differentiation, signaling, pigment production and secretion, and dendricity (Yuan and Jin, 2018).

The KC:MC unit acts as a first line of defense to respond to environmental stimuli including ultraviolet (UV) light from the sun. Melanin transfer to KCs is a critical protective response that helps prevent UV-induced DNA damage and subsequent mutagenesis leading to cancer (Brenner and Hearing, 2008b; Yamaguchi *et al.*, 2006). Activation of the MC transcription factor MITF in response to secretion of KC-derived factors (e.g. α–MSH, KITL, ET1; Melanocyte stimulating hormone, Kit ligand, Endothelin 1) following UV exposure results in upregulation of melanin-producing enzymes (including TRP1; Tyrosinase related protein 1), and melanin production and secretion (Nguyen and Fisher, 2018; Serre *et al.*, 2018). UV exposure also initiates signaling cascades in both KCs and MCs, resulting in secretion of cytokines and chemokines including interleukins (IL1, 3, 6, 8), interferon (IFNγ), granulocyte-colony stimulating factor (G-CSF), and tumor necrosis factor (TNFα) (Schwarz and Luger, 1989; Terazawa and Imokawa, 2018; Yoshizumi *et al.*, 2008), all of which control multicellular reactions such as the tanning response within the KC:MC unit. Lengthened dendrites in response to UV light increase MC interactions with KCs to facilitate melanin transfer (Lopez *et al.*, 2015; Weiner *et al.*, 2014). Further, secreted factors from KCs function to increase DNA damage repair pathways in MCs, reducing the risk of oncogenic mutations (Kadekaro *et al.*, 2010; Swope and Abdel-Malek, 2016).

While UV light can modulate expression of cell-cell adhesion proteins (Dusek *et al.*, 2006; Jamal and Schneider, 2002; Johnson *et al.*, 2014), the signaling roles of cadherin proteins have not been fully explored. In particular, the UV-sensitive (Johnson *et al.*, 2014), KC-specific desmosomal cadherin desmoglein 1 (Dsg1) has been linked to both cell adhesion and signaling as well as regulation of cytokines under homeostatic and pathologic conditions (Hammers and Stanley, 2013; Samuelov *et al.*, 2013). Outside of its adhesive function, Dsg1 is an important regulator of cellular signaling, specifically in attenuating the MAPK/ERK pathway to promote KC differentiation (Getsios *et al.*, 2009; Harmon *et al.*, 2013). Activation of the MAPK/ERK pathway increases epidermal inflammation following UV exposure (El-Abaseri *et al.*, 2013). Mutations in Dsg1 lead to Severe dermatitis, multiple Allergies, and Metabolic wasting (SAM) syndrome, accompanied by increased pro-allergy cytokines (Samuelov *et al.*, 2013). Also, several pathological conditions which result in downregulation of Dsg1 at the cell surface can induce cytokine production, suggesting a role for Dsg1 in this process (Hammers and Stanley, 2013; Ishii *et al.*, 2001; Polivka *et al.*, 2018; Sherrill *et al.*, 2014). Based on these previous studies, we questioned whether reduction of Dsg1, such as occurs following UV exposure or other environmental stimuli, could mediate changes in paracrine communication between KCs and MCs known to occur in response to these stimuli.

In the present study, we queried whether KC-specific Dsg1 can modulate the epidermal microenvironment and MC behavior through regulation of cytokines, chemokines, and other secreted factors, thereby contributing to the epidermal tanning response. Using conditioned media from KCs with altered Dsg1 expression, we tested effects of Dsg1 suppression and expression upon MC signaling, pigment secretion, and dendricity. An increase in paracrine signaling from KCs resulted in increased pigment secretion and altered dendrite length in MCs. We also utilized 3D human skin equivalents containing both KCs and MCs to study the role of Dsg1 reduction in MC localization within the epidermis, finding that suppression of Dsg1 resulted in mislocalization of MCs into the suprabasal layers. This study elucidates a new role for Dsg1 in regulating KC:MC cell-cell communication, and suggests a specific mechanism by which the epidermis senses and propagates signals in response to environmental stimuli, which when perturbed can result in MC dysplasia.

## RESULTS

### Suppression of KC Dsg1 increases cytokine production and secretion

We first sought to determine which cytokines were altered upon Dsg1 suppression in KCs. KCs infected with retrovirus expressing non-target small hairpin RNA (NTshRNA), Dsg1-shRNA, or Dsg1-shRNA reconstituted with silencing resistant Dsg1-Flag (Dsg1FL) were grown to confluence, switched to high calcium medium to induce differentiation, and assessed by qRT-PCR after 3 days. Cytokine transcripts including IL1α, IL1β, IL6, IL8, and CXCL1 were significantly upregulated, IL2, IL4, and IL10 were significantly downregulated, and IL17, IL19, IL23, TNFα and IFNγ were highly variable or not significantly altered by suppression of Dsg1 (Figure 1A, additional targets in Table 1). When a subset of these targets were tested, we found them to be returned to control levels upon restoration of Dsg1FL (Figure 1A). Depletion of another desmosomal cadherin in KCs, Dsg3, did not result in significant changes in tested cytokine mRNA levels (Figure 1B).

**Table 1.** Average fold change in KC cytokine transcripts upon depletion of Dsg1 (shCon with shDsg1) to accompany Figure 1A. IL, Interleukin, CXCL1, Chemokine ligand 1; TNF, Tumor Necrosis Factor; IFN, Interferon. Tested by qRT-PCR (N=3 *p<0.03).

| <u>Target</u> | <u>Avg. Fold Change</u> | <u>p-value</u> |
| --- | --- | --- |
| IL2 | 0.2709 | *0.0314 |
| IL4 | 0.3416 | *0.0105 |
| IL10 | 0.4813 | *0.0090 |
| IL17 | 2555.6718 | 0.1213 |
| IL19 | 58.6191 | 0.1191 |
| IL23 | 40.7489 | 0.0791 |
| CXCL1 | 2.3038 | *0.0149 |
| TNFα | 22.9530 | 0.0816 |
| IFNγ | 43.0594 | 0.0686 |

**Figure 1.**
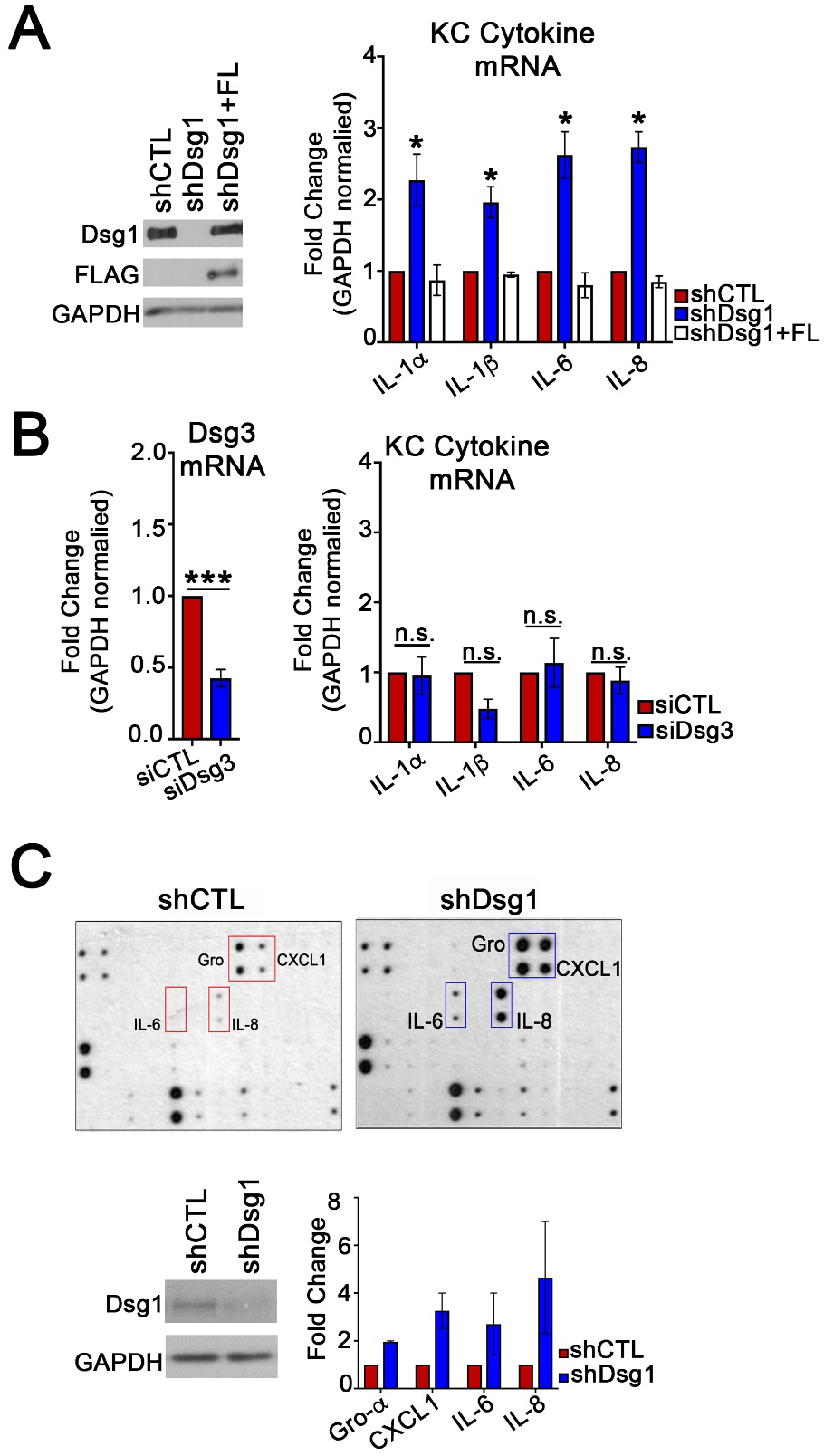
Suppression of KC desmoglein 1 increases cytokine production and secretion. (**A**) NTshRNA (shCTL), Dsg1shRNA (shDsg1), and shDsg1 plus silencing resistant Dsg1-Flag (FL)-infected KCs were induced to differentiate by switching from 0.07 to 1.2 mM CaCl_2_-containing medium for 72 hours. The mRNA was collected, and cDNA was analyzed for transcription of several cytokines (see Table 1 for additional tested cytokines) (N=3; *p<0.03). (**B**) Cells with siRNA targeting Dsg3 were also analyzed and found not to have significantly changed levels of cytokines compared to controls (N=3; n.s. = not significant, *p<0.03, ***p<0.0002). (**C**) Media conditioned from NTshRNA and Dsg1shRNA-infected KCs from days 3-5 after being inducted to differentiate was incubated with a Human Cytokine C3 Array (Raybio) to detect secretion of up to 42 cytokines, chemokines, and growth factors. Secretion of GRO, CXCL1, IL6, and IL8 were increased upon suppression of Dsg1 compared to controls (N=2).

To determine which of these altered cytokines were being secreted from KCs, media conditioned days 3-5 (a 48 hour time period) after initiation of differentiation by growth in high calcium medium was collected from KCs infected with NTshRNA or Dsg1shRNA. Dot-blot analysis of 42 targets revealed increased secretion of GRO, CXCL1, IL6, and IL8 in the Dsg1shRNA conditioned media compared to NTshRNA conditioned media (Figure 1C). KCs exposed to UV light have been reported to increase secretion of IL6 and IL8 among other cytokines/chemokines (Schwarz and Luger, 1989; Yoshizumi *et al.*, 2008). Since Dsg1 is downregulated in KCs following UV exposure (Johnson *et al.*, 2014) and its suppression results in increased cytokine/chemokine secretion, we hypothesized that Dsg1 may play a role in coordinating the skin’s paracrine response to environmental insults.

### Exposure to Dsg1-deficient KC conditioned media increases MC pigment secretion through classic ligand-dependent mechanisms

Since induction of pigment production and secretion in MCs is a crucial protective response in the epidermis that can be regulated by paracrine signaling (Serre *et al.*, 2018; Yuan and Jin, 2018), we next tested MC pigment secretion following incubation with conditioned media from KCs with normal or suppressed Dsg1 expression. MCs were grown for 7 days in a 1:1 mixture of MC media and media conditioned by KCs days 3-5 after initiation of differentiation. Fresh 1:1 mixture was added to the culture every 2 days (none of the previously depleted media was removed). On day 7, media was harvested and melanin secretion was analyzed by testing absorbance at 405nm (Laskin *et al.*, 1982). Conditioned media from Dsg1-deficient KCs increased MC pigment secretion compared to control media or media from KCs with restored Dsg1FL (Figure 2A). As shown in the images below the graph (Figure 2A), there was notable clonal variability in coloration of pigment secretion, but all tested MC clones responded to incubation with Dsg1-deficient conditioned media by increasing pigment secretion (Compiled data presented in Supplemental Figure 1).

**Figure 2.**
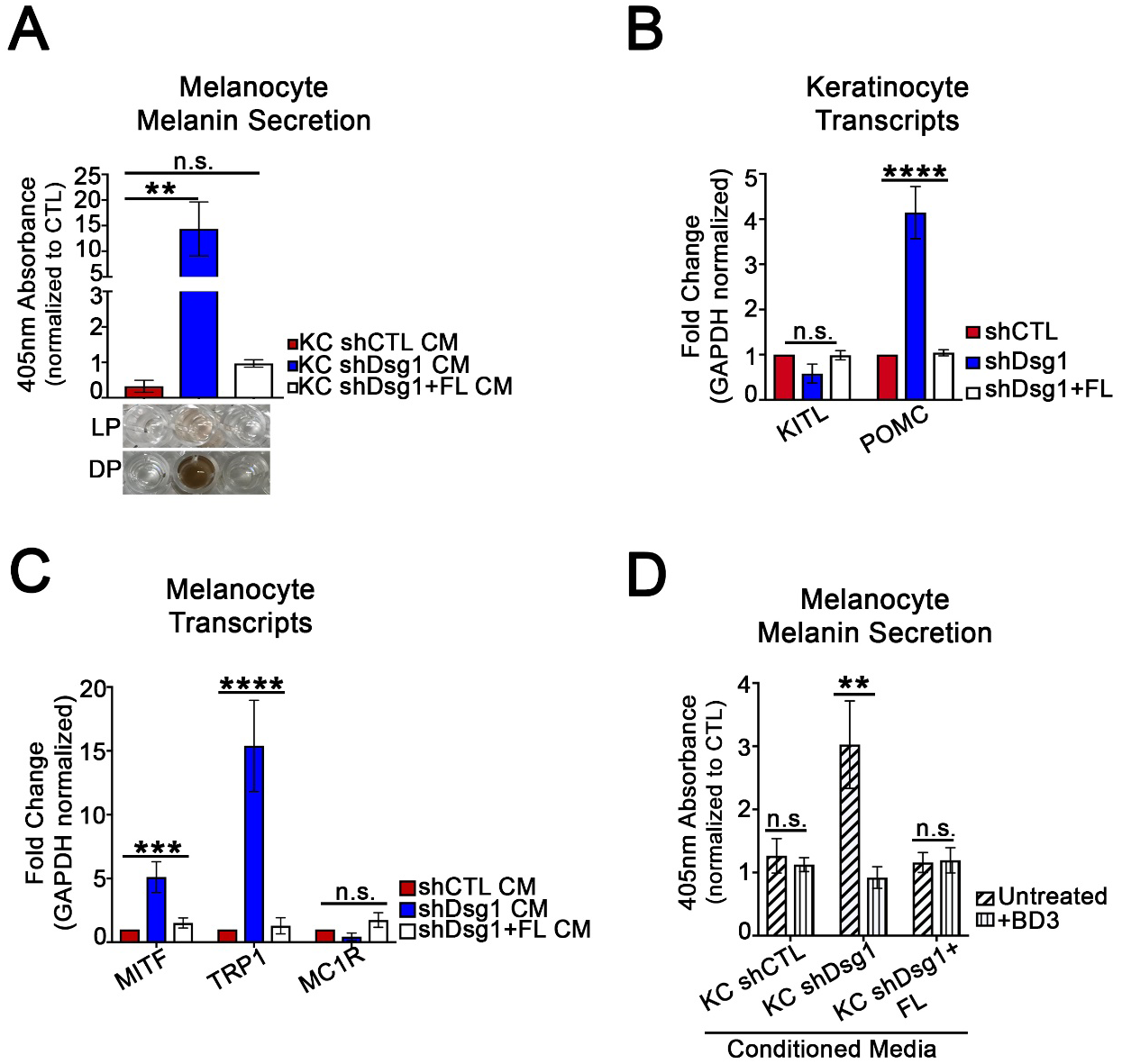
Exposure to Dsg1-deficient KC conditioned media increases MC pigment secretion through classic ligand-dependent mechanisms. **(A)** Conditioned media (CM) from Dsg1-deficient (shDsg1) KCs increased MC pigment secretion compared to control (CTL) media or media from KCs expressing exogenous full length Dsg1 (Dsg1FL) (Graph: CM from KC clones were tested on 1 MC clone; n.s. = not significant, **p<0.002). Multiple clones of MCs were tested with multiple sets of conditioned media (Supplemental Figure 1). Pigment secretion from two MC clones (LP = lightly pigmented, DP = darkly pigmented MC donor) are pictured to indicate the clonal variation but consistent phenomenon. **(B)** *Pro-opiomelanocortin* (POMC) mRNA is increased in shDsg1 KCs, while KIT ligand is not changed (N=3; ****p<0.0001). **(C)** Transcripts involved in pigment production in MCs (Melanogenesis associated transcription factor [MITF], Tyrosinase-related protein 1 [TRP1], and Melanocortin 1 receptor [MC1R]) are upregulated following incubation of MCs with CM from Dsg1 deficient KCs (N=3; ***p<0.0002, ****p<0.0001). **(D)** Inhibiting the MC Melanocortin 1 receptor (MC1R) using recombinant human Beta defensin 3 (BD3) in KC CM inhibits MC pigment secretion (N=3; **p<0.002).

To determine whether classical inducers of pigment production and secretion were induced in KCs upon Dsg1 silencing, we tested for mRNA expression levels of *Pro-opiomelanocortin* (*POMC* - precursor for the melanocortin 1 receptor [MC1R] ligand α-MSH), *KITL*, and *ET1*. *POMC* was increased upon Dsg1 suppression in KCs while *KITL* was not (Figure 2B). *ET1* was highly variable (data not shown). We tested induction of *Mitf* and *Trp1* in MCs treated with conditioned media, finding both to be elevated in MCs incubated with conditioned media deficient in Dsg1 (Figure 2C). When MCs were incubated with human beta defensin 3 (BD3) to inhibit MC1R (Swope *et al.*, 2012; Wolf Horrell *et al.*, 2016), pigment induction was inhibited downstream of Dsg1-deficient conditioned media (Figure 2D). Together these results indicate that reduction of Dsg1 in KCs results in induction of the classical pigment production pathways in MCs dependent upon paracrine signaling from KCs.

### Dsg1-deficient KCs change MC dendricity, partially dependent upon cytokine/chemokine signaling

MC dendrites lengthen and branch in response to UV light, increasing MC interactions with KCs to facilitate melanin transfer (Lopez *et al.*, 2015; Weiner *et al.*, 2014). To determine contributions of KC Dsg1 reduction to MC morphological changes, MCs were incubated with conditioned media from NTshRNA or Dsg1shRNA for either 12 hours or 7 days. The 12 hour incubation of MCs with Dsg1-deficient KC conditioned media resulted in increased dendrite length as compared to those in NTshRNA or Dsg1FL-conditioned media (Figure 3A), similar to the response seen following UV exposure and an important determinant of the number of KCs that a MC can interact with in normal skin (Lopez *et al.*, 2015; Weiner *et al.*, 2014). However, 7 days of exposure to Dsg1-deficient KC conditioned media resulted in shortening of MC dendrites (Figure 3A lower).

**Figure 3.**
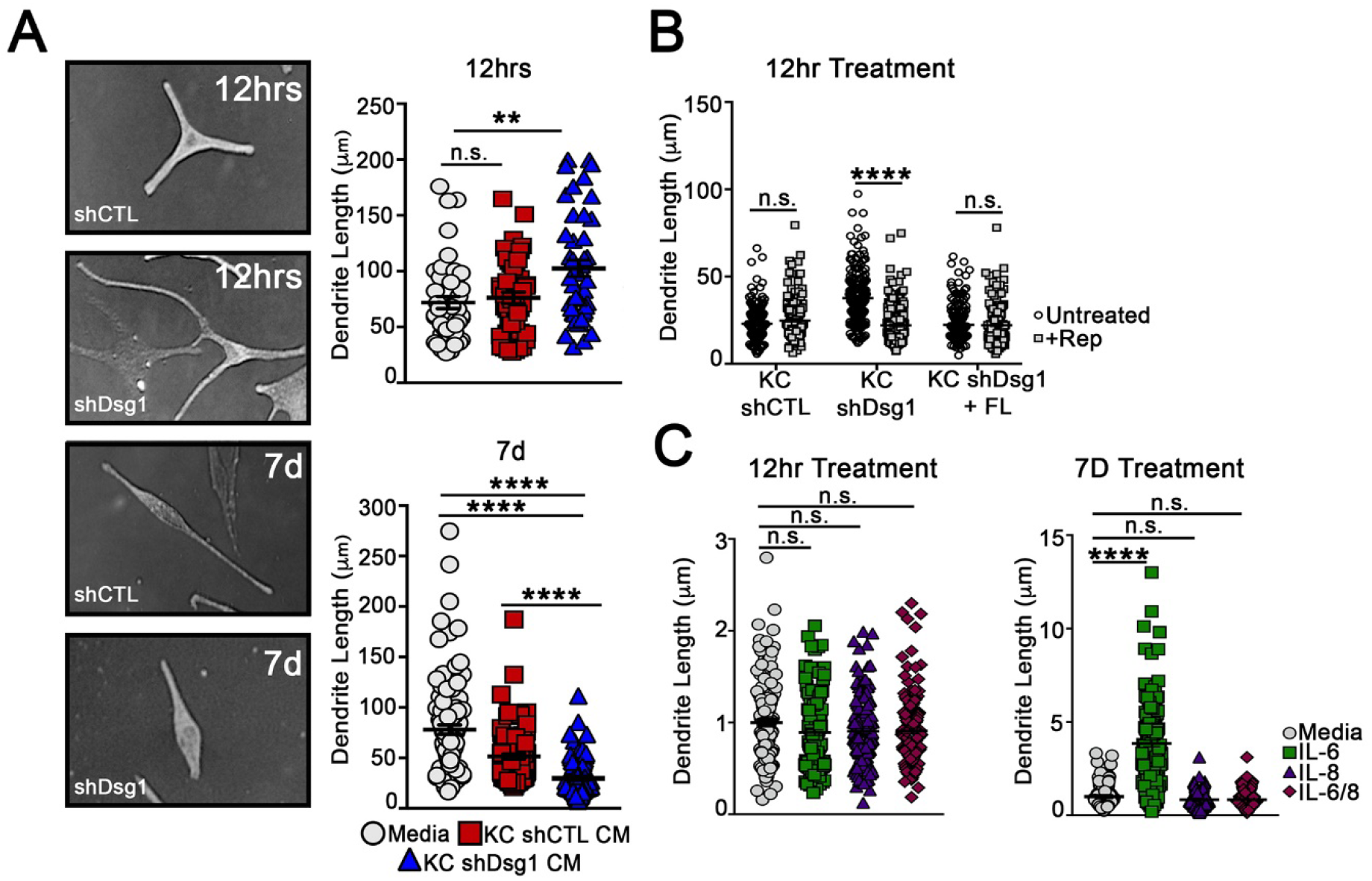
Dsg1-deficient KCs change MC dendricity, partially dependent upon cytokine/chemokine signaling. (**A**) MCs acutely exposed (12 hrs) to NTshRNA or Dsg1shRNA KC conditioned media exhibit increased dendrite length, while long-term exposure (7 days), results in dendrite shortening. (N=3 experimental replicates, >300 dendrites counted per condition; **p<0.002, ****p<0.0001). **(B)** Incubation of MCs with Reparixin, an inhibitor of the IL8/CXCL1 receptor CXCR2, inhibited the increase in MC dendrite length associated with acute exposure (12 hours) to Dsg1shRNA KC conditioned media (N=3 experimental replicates, >150 dendrites counted per condition; ****p<0.0001). **(C)** Incubation of MCs with IL6, IL8, or IL6 +IL8 in base MC media for 12 hours or 7 days. IL6 significantly increased MC dendrite length but only at the 7 day time point, while IL8 did not have a significant impact on dendrite length (N=3 experimental replicates, >150 dendrites counted per condition; ****p<0.0001).

We tested whether the cytokines secreted from KCs upon Dsg1 suppression were involved in the observed changes in MC dendricity. We inhibited the CXCL1/IL8 receptor (CXCR2) using Reparixin, which inhibited the increase in MC dendrite length at the 12 hour time point upon exposure to Dsg1-deficient conditioned media (Figure 3B). We also queried whether recombinant IL6, IL8, or a combination of the 2 added into base MC media for either 12 hours or 7 days were sufficient to impact dendrite length. While neither individual cytokine was sufficient to alter dendrite length at the 12 hour time point, MC dendrites significantly lengthened in the presence of IL6, but not IL8 after incubation for 7 days (Figure 3C). This observation is consistent with the possibility that individual cytokines contribute to MC dendrite length changes, but are not sufficient to recapitulate the time course and/or signal propagation stimulated by the complete Dsg1-deficient conditioned media. This may indicate that multiple paracrine factors are required to elicit the MC dendrite changes observed upon addition of Dsg1-deficient conditioned media. Together these data indicate that cytokine/chemokine signaling is at least partially responsible for altering MC dendrite length downstream of Dsg1 suppression.

### MCs are mislocalized within the 3D skin structure when KC Dsg1 is reduced

The 3D architecture of the skin is an important contributor to how signals are propagated throughout the tissue and represents a complex integration of multiple signaling and adhesive cues that differ from when cells are grown in 2D culture (Li *et al.*, 2011). We assessed whether reduction of KC Dsg1 in 3D organotypic cultures containing both KCs and MCs would result in changes in MC behavior. Either NTshRNA or Dsg1shRNA-infected KCs were seeded with MCs at a physiological ratio of 36:1, grown, and lifted to the air-liquid interface to allow stratification for six days. MCs were mis-localized from the basal to suprabasal layers in KC Dsg1-deficient organotypic cultures (Figure 4A). Imaging of organotypic cultures (shCon, shDsg1; representative immunoblot Figure 4B) prepared using the whole mount method revealed the extent to which melanocytes moved into the upper epidermal layers throughout the entire organotypic culture rather than in one plane of section (Figure 4C).

**Figure 4.**
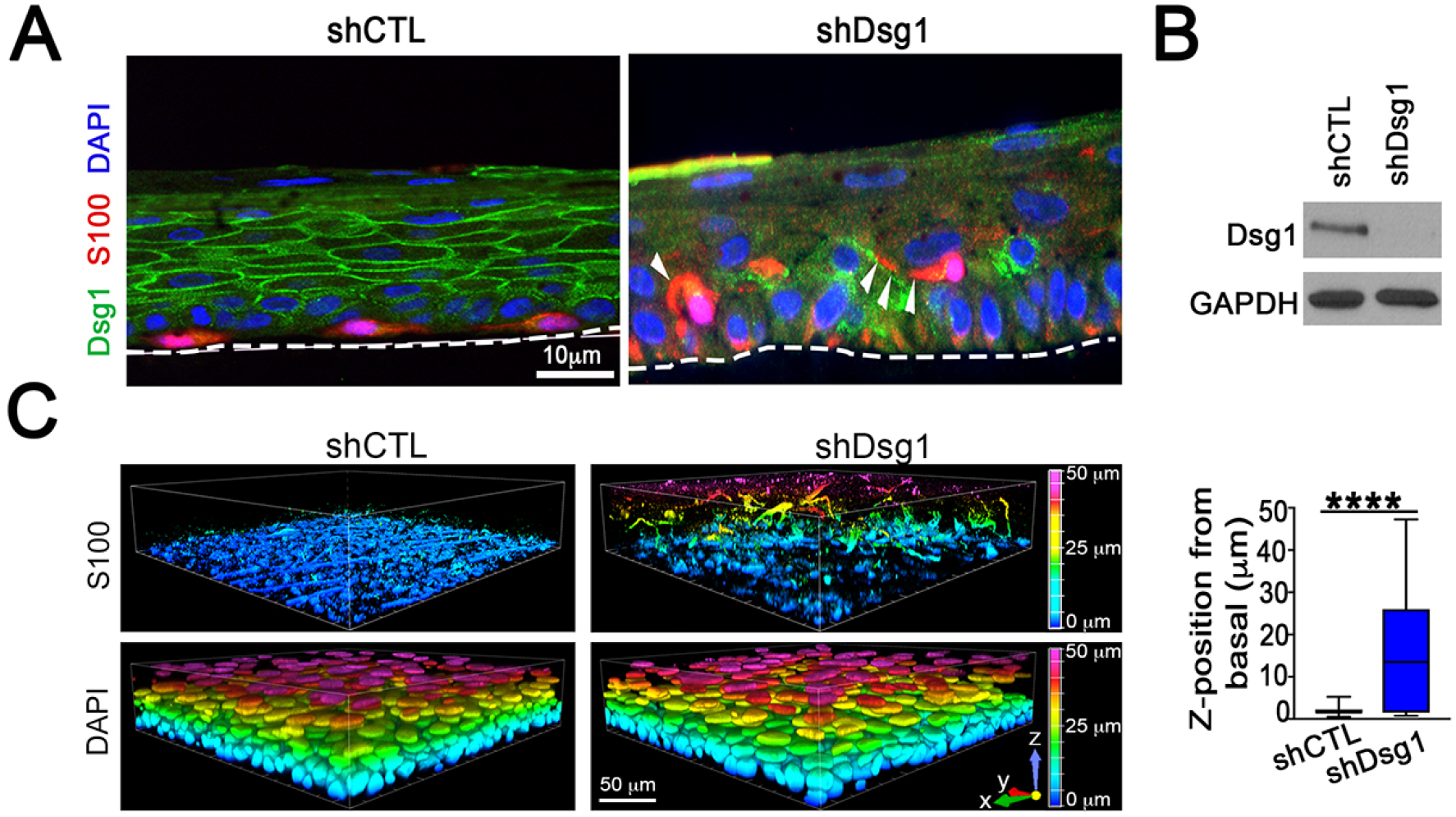
MCs are mislocalized within the 3D skin structure when KC Dsg1 is reduced. **(A)** Silencing Dsg1 in KCs co-cultured with MCs in 3D reconstructed human epidermis results in mislocalization of MCs from basal to suprabasal layers. Dsg1 marks KCs and can be seen to be mostly absent in the shDsg1 culture, S100 marks MCs, DAPI marks nuclei. White arrows indicate MC dendrites. Scale bar = 10 µm. **(B)** Immunoblot of Dsg1 protein expression for organotypic cultures pictured in C. GAPDH is a loading control. **(C)** Images of 3D organotypic cultures of KCs and MCs in presence and absence of Dsg1 prepared using the whole mount method. S100 marks MCs, showing their movement into the suprabasal layers upon Dsg1 reduction (N=3; ****p<0.0001). Horizontal scale bar = 50 µm, vertical color scale bar = Z position. Graphical representation is of MC Z position from basal layer.

### Dsg1 levels are significantly reduced at cell-cell borders in the KCs surrounding human dysplastic nevi and melanoma

The observed changes in MC dendrite length, localization, and cytokine profiles when exposed to conditioned media from KCs with suppressed Dsg1 are reminiscent of MCs that have undergone transformation (Colebatch and Scolyer, 2018; Dhawan and Richmond, 2002). Increased MITF levels have also been associated with MC transformation and melanoma progression (Hartman and Czyz, 2015). We therefore examined Dsg1 levels in KCs surrounding benign and dysplastic pigmented nevi and in early stage melanomas to determine if Dsg1 suppression was a phenomenon observable in the progression of human disease. To assess the global levels of Dsg1 at KC cell-cell borders in normal human skin and melanoma samples, tissue sections were stained for Dsg1 and the intensity measured. Control skin and benign MC nevi exhibited similar levels of Dsg1 at cell-cell borders (Figure 5A, 5B upper). However, Dsg1 was found to be significantly decreased in matched dysplastic nevi and melanoma samples from the same individuals compared to normal skin. Epithelial cadherin, (Ecad) was not significantly reduced in the same KC populations, indicating that reduction in Dsg1 precedes the previously reported reduction in Ecad in human melanoma (Figure 5A, 5B lower) (Lee and Herlyn, 2007). Together our data are consistent with the idea that decreased KC Dsg1 levels as occurs following UV exposure (Johnson *et al.*, 2014) alter the microenvironment within the KC:MC unit to propagate the tanning response. We posit that longer-term Dsg1 suppression, as observed surrounding human dysplastic and melanoma lesions may contribute to melanocyte transformation.

**Figure 5.**
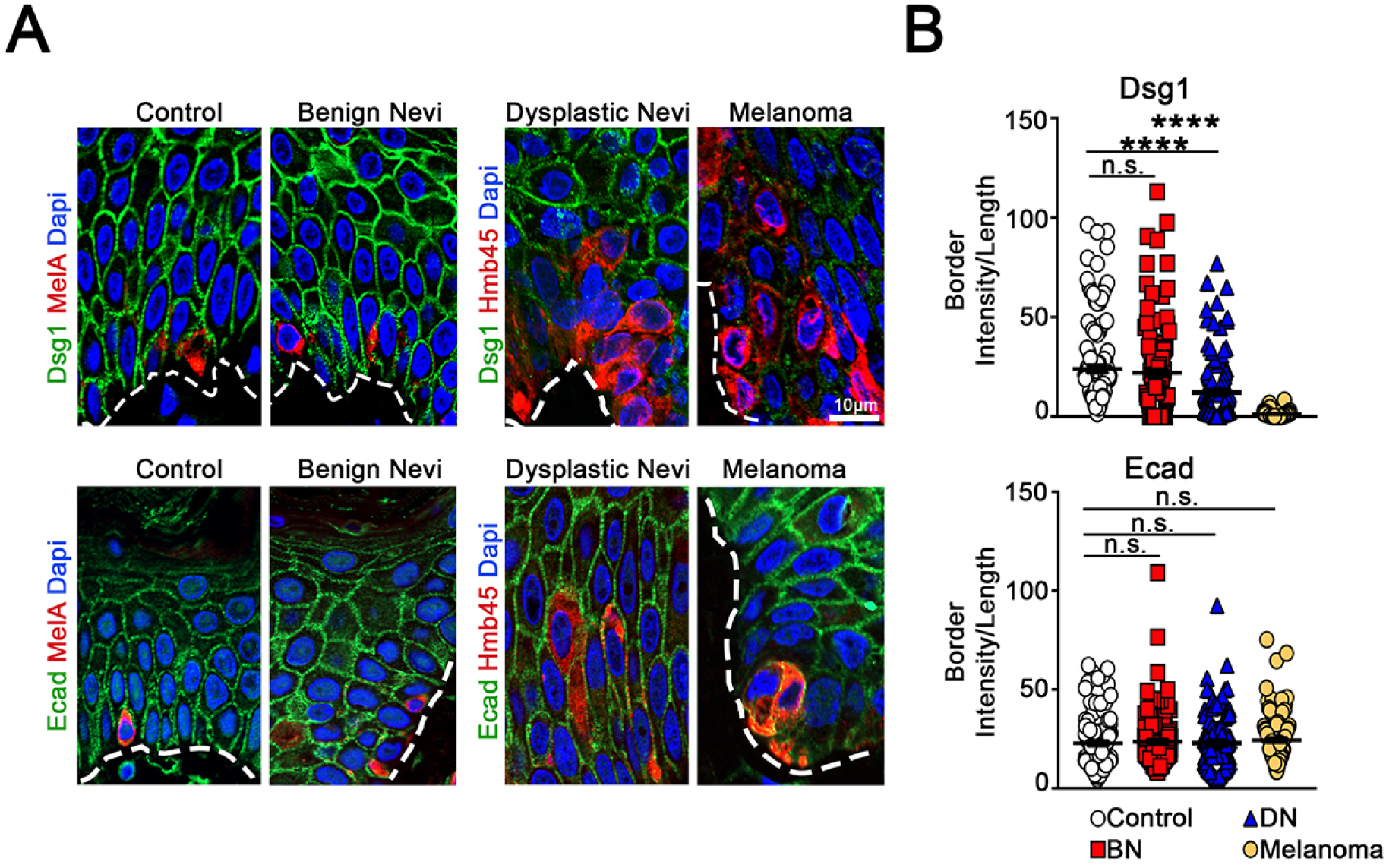
Dsg1 levels are significantly reduced at cell-cell borders in the KCs surrounding human dysplastic nevi and melanoma. (**A**) Staining of Dsg1, Ecad, and the MC markers MelA in control and benign nevi and HMB45 in matched dysplastic and melanoma tissues from the same individuals. (**B**) Quantification of Dsg1 and Ecad border intensity as compared to control tissue. BN = Benign Nevi, DN = Dysplastic Nevi. Scale bar = 10 µm. N=12/condition (120 borders/condition). ****p<0.0001.

## DISCUSSION

Our data support a model where Dsg1, important in epidermal cell-cell adhesion, also functions as a rheostat to sense and respond to environmental insults; reduced Dsg1 expression such as occurs following UV light exposure (Johnson *et al.*, 2014) increases paracrine signaling from KCs to help initiate the MC tanning response (Figure 6). The regulation of MCs via secretion of paracrine factors by KCs is a well-established phenomenon (Serre *et al.*, 2018; Yuan and Jin, 2018). Our study places Dsg1 as a key regulator of the microenvironment within the KC:MC unit.

**Figure 6.**
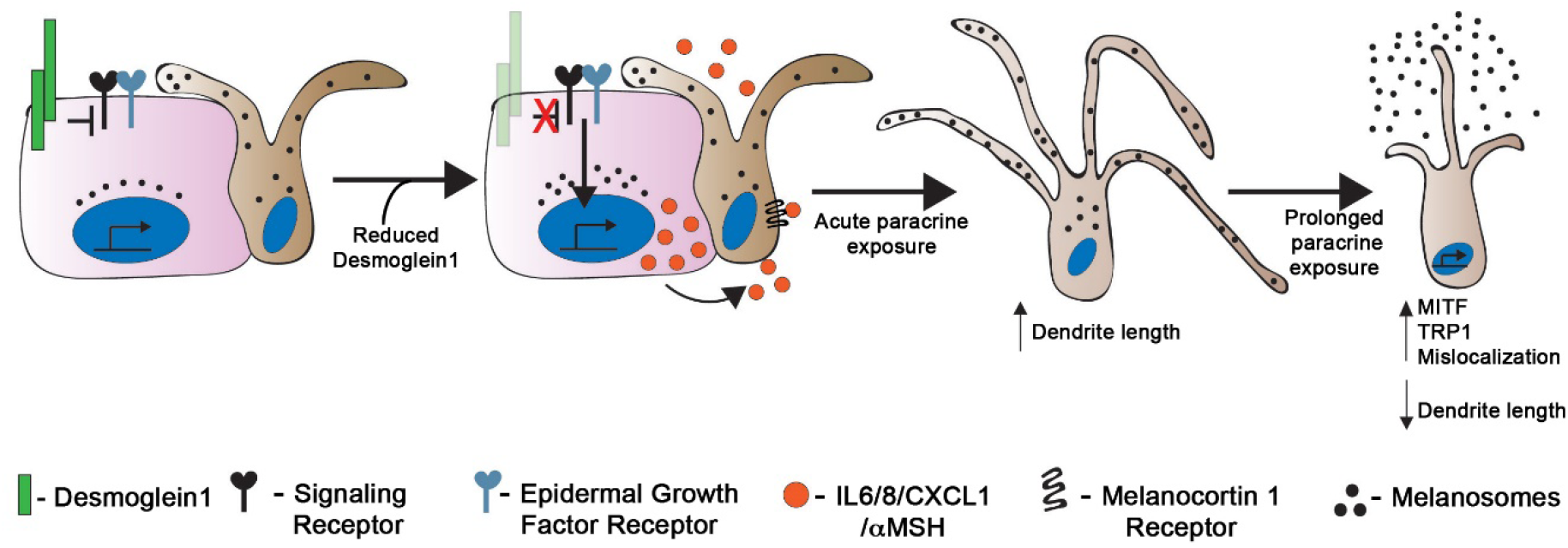
Model: Keratinocyte desmoglein 1 regulates the epidermal microenvironment and melanocyte behavior. Keratinocytes (pink) and melanocytes (tan) in the skin communicate with each other through both direct contact and paracrine signaling. Reduction in expression of keratinocyte Dsg1 initiates a signaling cascade that results in upregulation and secretion of paracrine factors including IL6, IL8, CXCL1, and αMSH that affect the signaling, morphology, pigment secretion, and localization of neighboring melanocytes.

Protection against damage from environmental exposure, including UV light from the sun is one of the key barrier functions of the epidermis. The classic KC:MC tanning response protects against mutagenesis (Brenner and Hearing, 2008a; Kadekaro *et al.*, 2010; Swope and Abdel-Malek, 2016; Yamaguchi *et al.*, 2006) and involves KC increase of POMC, production and secretion of αMSH as well as other secreted factors, and subsequent activation of the MC receptor MC1R, downstream activation of MITF, and pigment production through TRP1 and TRP2 (Serre *et al.*, 2018). All human skin, regardless of skin type or melanin type and quantity (Wakamatsu *et al.*, 2006) undergoes reaction to UV exposure including increased pigmentation (Alaluf *et al.*, 2001). A recent study suggested that MCs from darkly pigmented individuals more effectively increased melanogenesis after UVB exposure in the presence of KC conditioned media compared to MCs from lightly pigmented individuals, suggesting an enhanced photoprotective mechanism in darker skin types (Lopez *et al.*, 2015). Though a full study of MC clones from light and darkly pigmented individuals was not undertaken here, we show that reduction in KC Dsg1, such as occurs following UVB light exposure (Johnson *et al.*, 2014) increases pigment secretion in MCs from donors with multiple skin types.

MC dendrites are crucial for mediating the tanning response following UV exposure since they are the vehicle through which melanosomes are transferred to multiple KCs (Gilchrest *et al.*, 1996). We showed that factors present in Dsg1-deficient conditioned media can signal to MCs to alter dendrite length. UV exposed KCs increase secretion of IL6 and IL8, similar to our findings of cytokine secretion downstream of Dsg1 suppression (Schwarz and Luger, 1989; Yoshizumi *et al.*, 2008). We showed that IL6 and the IL8/CXCL1 receptor (CXCR2) both impact melanocyte dendricity downstream of Dsg1. However, addition of IL6 or inhibition of CXCR2 were both insufficient to recapitulate the precise effects and timeline of complete Dsg1-deficient KC media. Therefore, further studies are needed to elucidate the roles of these and other cytokines and chemokines in regulating the dendrite morphology of MCs downstream of Dsg1.

MCs were mislocalized away from the basement membrane when grown in the context of KCs depleted of Dsg1 in 3D epidermal skin equivalents, reminiscent of the pagetoid appearance of MCs in dysplastic nevi or in melanoma in situ (Colebatch and Scolyer, 2018). It is possible that repeated cycles of acute UV exposure could reduce Dsg1 expression in KCs for prolonged amounts of time, leading to alternations in other KC:MC adhesive molecules (Mescher *et al.*, 2017) or an altered microenvironment within the KC:MC unit to promote MC dysplasia.

Nests of KCs surrounding human dysplastic and early stage melanoma lesions exhibited decreased levels of Dsg1 at cell-cell borders which preceded the previously reported loss of Ecad during melanoma progression (Lee and Herlyn, 2007). We show here that normal KCs lacking Dsg1 increase secretion of IL6, IL8, and CXCL1, all of which have been implicated in melanoma development and maintenance (Jobe *et al.*, 2016; Payne and Cornelius, 2002). While our study implicates Dsg1 loss in regulation of the protective epidermal tanning response, the possibility exists that repeated reduction in KC Dsg1, such as through repeated acute UV exposure or through an increased epidermal oxidative environment (Wittgen and van Kempen, 2007), ultimately leads to a pro-tumorigenic microenvironment in vivo. Transformed MCs themselves could secrete factors resulting in more sustained loss of KC Dsg1. Cytokine secretion and altered inflammatory microenvironment contributes to the growth, invasion, and angiogenesis of transformed MCs and melanoma cells (Richmond *et al.*, 2009). In fact, CXCL1 was identified as one of the most highly modified markers allowing discrimination between common melanocytic and dysplastic nevi (Mitsui *et al.*, 2016). Identifying KC Dsg1 as a regulator of MC pigment secretion, dendricity, and position within epidermis places Dsg1 at a nexus of regulating normal homeostasis within the KC:MC unit, and suggests how this can go awry in nevi to melanoma progression as MCs themselves begin to alter the microenvironment, questions which have remained open in the field for decades and remain open still (Colebatch and Scolyer, 2018; Haass and Herlyn, 2005).

Modulating KC Dsg1 levels through topical formulations could alter the epidermal microenvironment and cross-talk within the KC:MC unit. Histone deacetylase (HDAC) inhibition was shown to moderately increase KC Dsg1 levels after UVB exposure (Johnson *et al.*, 2014) and HDAC inhibitors are being formulated for topical use in management of cutaneous diseases (Fournier *et al.*, 2018). Other desmogleins are being regulated through use of glucocorticoids, rapamycin, and Stat3 inhibition in treatment of pemphigus (Mao *et al.*, 2017). Therefore, pharmacological modulation of Dsg1 expression is possible with potential ramifications for clinically managing pigmentation or reducing melanomagenesis from dysplastic nevi.

In conclusion, this study demonstrates a new role for KC Dsg1 in regulating paracrine cross-talk within the epidermal KC:MC unit. Reduction in Dsg1 results in KCs increasing both ligand-dependent and cytokine/chemokine-dependent signaling to MCs to stimulate MC pigment production and secretion, and dendricity. Within the 3D epidermis, the presence of KC Dsg1 helps properly position MCs in the basal layer; Dsg1 reduction results in suprabasal mislocalization of MCs, which may be involved in the trajectory from benign to dysplastic nevi formation, since KC Dsg1 loss occurs surrounding human dysplastic nevi and melanoma lesions.

## MATERIALS AND METHODS

### Cell culture and retroviral transduction

KCs and MCs were isolated from neonatal foreskin provided by the Northwestern University Skin Disease Research Center (NUSDRC) as described in (Halbert *et al.*, 1992). KCs were propagated in M154 medium supplemented with human KC growth supplement (Life Technologies, Grand Island, NY), 1,000 x gentamycin/amphotericin B solution (Life Technologies, and 0.07mM CaCl_2_ (low calcium). Confluent KC monolayers were induced to differentiate by the addition of 1.2 mM CaCl_2_ (high calcium) in M154 in the presence of human KC growth supplement. KCs were transduced with retroviral supernatants produced from Phoenix cells (provided by G. Nolan, Stanford University, Stanford, CA) as previously described (Getsios *et al.*, 2004; Simpson *et al.*, 2010b). MCs were cultured in OptiMEM (Life Technologies, Carlsbad, CA) containing 1% penicillin/streptomycin (Corning, Corning, NY), 5% fetal bovine serum (Sigma-Aldrich, St. Louis, MO), 10 ng/ml bFGF (ConnStem Inc., Cheshire, CT), 1 ng/ml heparin (Sigma-Aldrich), 0.1 mM N^6^, 2’-O-dibutyryladenosine 3:5-cyclic monophosphate (dbcAMP; Sigma-Aldrich), and 0.1 mM 3-isobutyl-1-methyl xanthine (IBMX; Sigma-Aldrich).

### Organotypic skin cultures and whole mount samples

Organotypic cultures were grown as described previously (Arnette *et al.*, 2016; Getsios *et al.*, 2009) with the addition of the following steps: the collagen plug was coated with extracellular matrix using 804G supernatant (Langhofer *et al.*, 1993). MCs were seeded onto the collagen plug overnight. KCs were then seeded at a physiologic ratio of approximately 36 KCs:1 MC. Organotypic cultures fixed in 10% neutral buffered formalin were embedded in paraffin blocks and cut into 4 µm sections. For indirect immunofluorescence microscopy, slides were baked at 60° C, de-paraffinized by xylenes, dehydrated with ethanol, rehydrated in PBS and permeablized by 0.5% Triton X-100 in PBS. Antigen retrieval was performed by incubation in 0.01 M Citrate buffer (pH 6.0) at 95°C for 15 minutes. Sections were blocked in 1% BSA/2% Normal Goat Serum/PBS for 30 minutes at 37°C. Primary antibody incubation was carried out overnight at 4°C in blocking buffer followed by washing in PBS. Secondary antibody incubation was carried out at 37°C for 45 minutes followed by washing in PBS. Sections were stained with 4′,6-Diamidino-2-phenylindole (DAPI - Sigma-Aldrich) at a final concentration of 5 ng/µl at room temperature for 2 minutes followed by washing in PBS and water. Cover slips were mounted on the sections with ProLong Gold Antifade Reagent (Invitrogen, Life Technologies).

For whole mount imaging: Six days after lifting to the air-liquid interface, the epidermal equivalent layer was removed from the collagen plug and fixed in 4% paraformaldehyde in PBS for 15 min on ice. Samples were then washed three times in PBS for 5 min each at room temperature.

Subsequently, samples were incubated in blocking buffer (1% Triton-X 100 with 5% goat serum in PBS) for 1 hr at 37°C followed by incubation overnight at 37°C with S100 (ab52642, anti-S100 beta; Abcam, Cambridge, UK) diluted at 1:100 in blocking buffer. Samples were washed 3 times for 10 min each with PBS at room temperature and then incubated overnight at 37°C with Alexa Fluor-conjugated secondary antibodies (1:250) in blocking buffer that included DAPI (2 µg/ml). Samples were washed 3 times for 10 min each with PBS at room temperature and mounted onto glass slides with Prolong Gold Antifade Reagent.

### Preparation of conditioned media from KCs

KCs infected at 20% confluence with LZRS-miR Dsg1, LZRS-NTshRNA, LZRS-Flag Dsg1, and LZRS-Dsg1Δ381 were grown to 80% confluence, expanded to 3 10 cm dishes, and grown to confluence. At confluence, cells were switched to high calcium medium and cultured for 3 days before refreshing the medium. On day 3 the medium was refreshed with new high calcium medium and harvested on day 5 (day 3-5 conditioned medium). This day 3-5 conditioned media (conditioned for a 48 hour time period) was used in all assays. All conditioned media was stored at - 20°C in aliquots until needed for an experiment.

### DNA constructs

LZRS-miR Dsg1 (Dsg1shRNA), LZRS-Flag Dsg1, and LZRS-Dsg1Δ381 were generated as described by (Getsios *et al.*, 2009) and (Simpson *et al.*, 2010a). LZRS – NTshRNA was generated as described in (Getsios *et al.*, 2004) with the following sequences inserted: NTshRNA1-fwd 5’ – GTATCTCTTCATAGCCTTAaa – 3’ and NTshRNA1-rev 5’ – ttTAAGGCTATGAAGAGATAC – 3’.

### Antibodies and reagents

The mouse monoclonal antibodies used were as follows: P124 (anti-Dsg1 extracellular domain; Progen, Heidelberg, Germany), 27B2 (anti-Dsg1 cytodomain; Life Technologies), and HMB45 (Melanoma gp100 antibody; Thermo Fisher Scientific). The rabbit monoclonal antibody EP1576Y (ab52642, anti-S100 beta; Abcam, Cambridge, UK) was used. Rabbit polyclonal antibodies used were: HECD1 (anti–E-cadherin; Takara, Kyoto, Japan), Flag (Cell Signaling Technologies), Anti-MelanA (ab15468), and GAPDH (G9545, glyceraldehyde-3-phosphate dehydrogenase; Sigma-Aldrich). Secondary antibodies for immunoblotting were goat anti-mouse and goat anti-rabbit peroxidase (Rockland; KPL, Gaithersburg, MD). Secondary antibodies for immunofluorescence were goat anti-mouse and goat-anti-rabbit linked to fluorophores of 488 and 568 nm (Alexa Fluor; Life Technologies). DAPI was used to stain nuclei.

### Western blot analysis of proteins

Whole cell lysates were collected from confluent monolayers in urea-SDS buffer (8 M urea/1% SDS/60 mM Tris (pH 6.8)/5% ß-mercaptoethanol/10% glycerol) and sonicated. Samples separated by SDS-PAGE were transferred to nitrocellulose, blocked in 5% milk/PBS, and incubated with primary antibodies in milk for 1 hour at room temperature or overnight at 4°C. After a series of PBS washes, secondary antibodies diluted in milk were added to blots. Protein bands were visualized using exposure to X-ray film. Densitometric analyses were performed of scanned immunoblots using ImageJ software.

### Dot Blot Analysis of Cytokines

Conditioned media from KCs (day 3-5 conditioned media) was concentrated 5x by centrifugation through a Centricon Plus-70 Centrifugal Filter with a cutoff of 3 kDa (Millipore Sigma, St Louis, MO) according to the manufacturer’s instructions. Protein concentration was normalized and a 1 mL sample was incubated according to the manufacturer’s instructions on a Human Cytokine C3 Array (Raybiotech, Norcross, GA) capable of detecting 42 human cytokines and chemokines.

### Quantitative real-time PCR

RNA was isolated from KCs and MCs using the RNeasy Mini Kit (Qiagen, Valencia, CA), according to the manufacturer’s instructions. MCs were grown in monoculture prior to being incubated overnight or for 7 days with either MC media alone or with a 1:1 mixture of MC media and conditioned media. Total RNA concentrations were equalized between samples and cDNA was prepared using the Superscript III First Strand Synthesis Kit (Life Technologies). Quantitative PCR was performed using SYBR Green PCR master mix (Life Technologies) and gene-specific primers (Table 2) in a StepOnePlus instrument. Calculations for relative mRNA levels were performed using the ΔΔCt method, normalized to GAPDH.

**Table 2.**
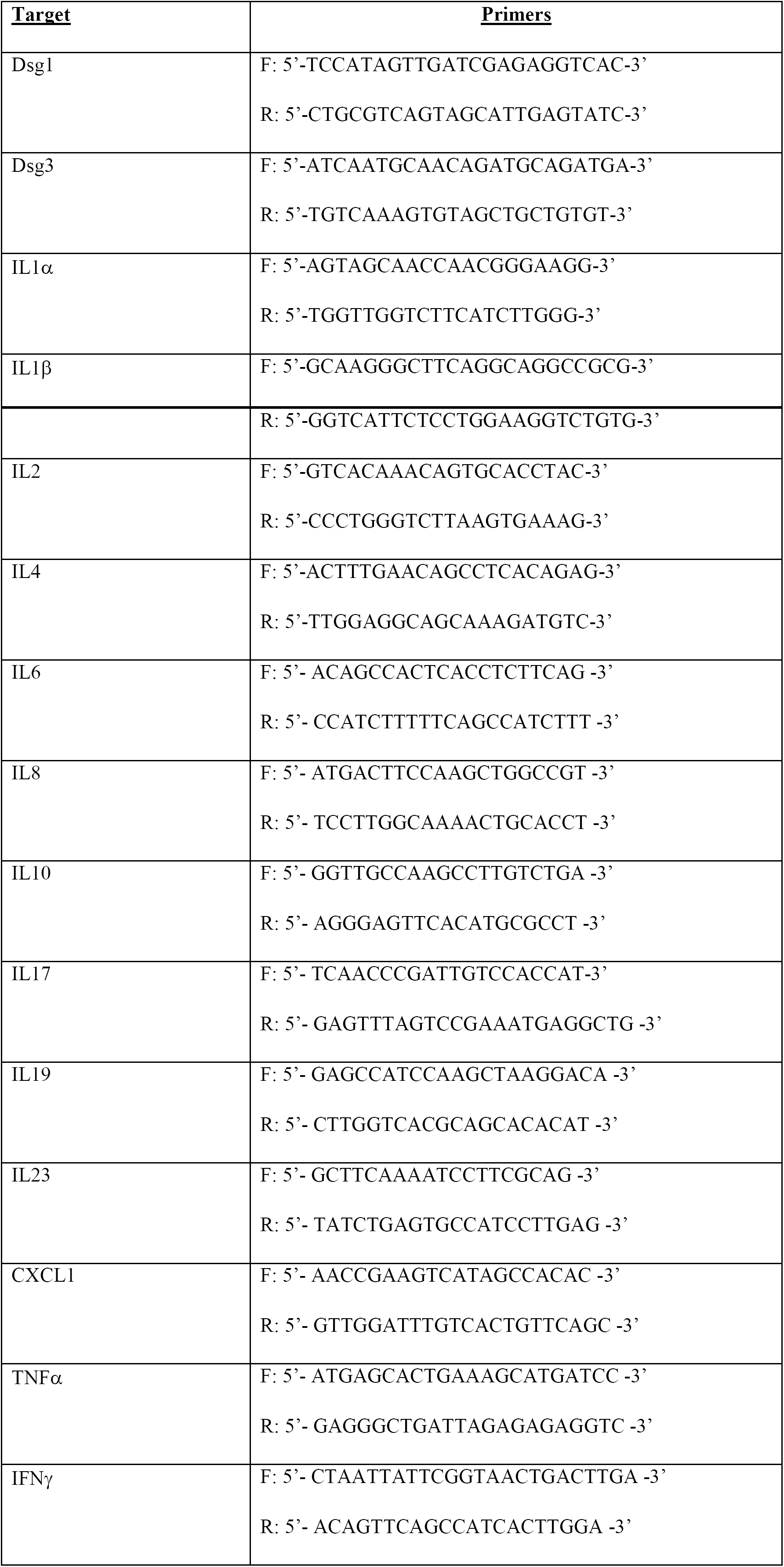
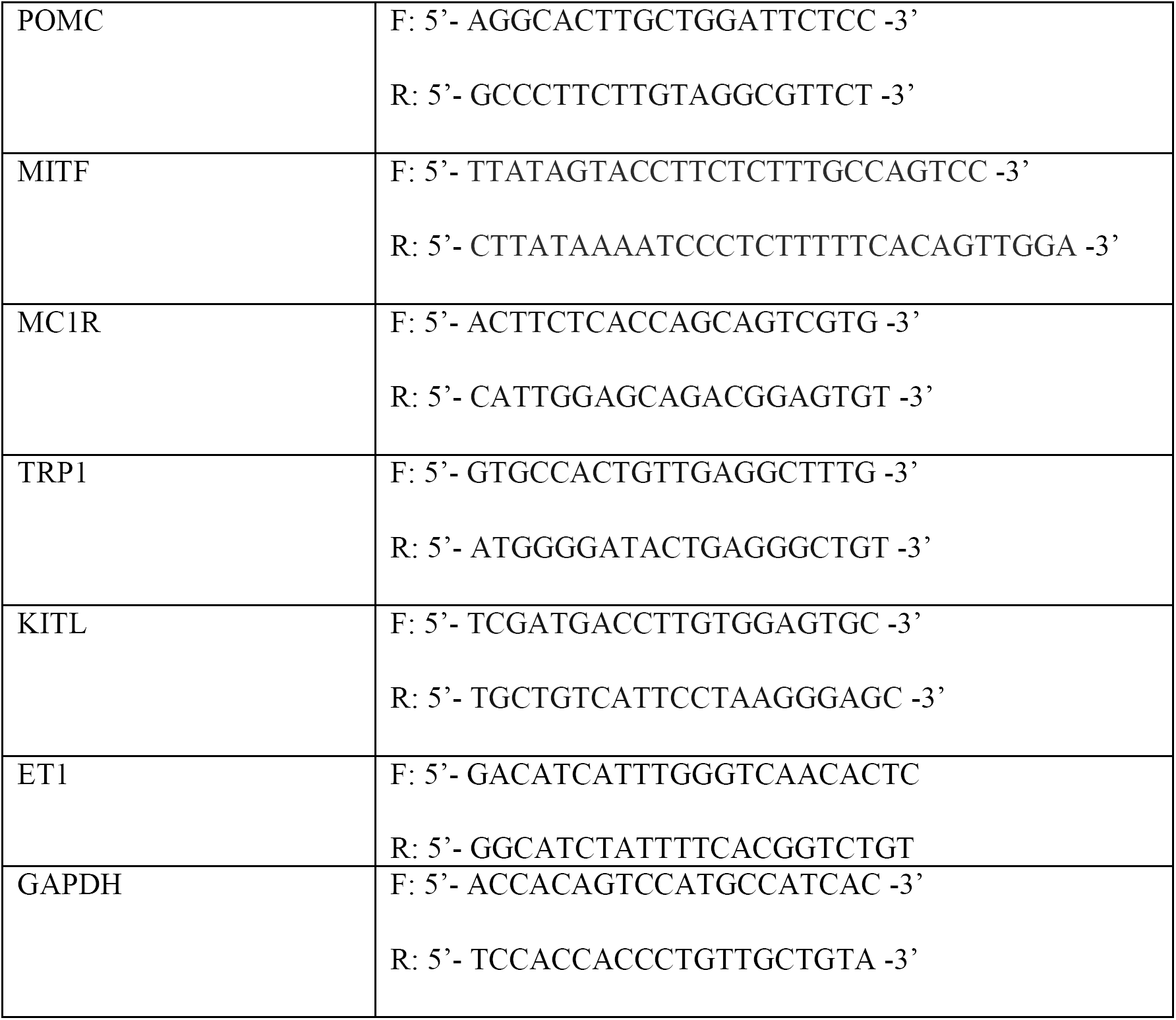
Primers used in this study. Dsg1, Desmoglein 1; Dsg3, Desmoglein 3; IL, Interleukin, CXCL1, Chemokine ligand 1; TNF, Tumor Necrosis Factor; IFN, Interferon; POMC, Pro-opiomelanocortin; MITF, Melanogenesis associated transcription factor; MC1R, Melanocortin 1 receptor; TRP1, Tyrosinase related protein 1; KITL, Kit ligand; ET1, Endothelin 1; GAPDH, Glyceraldehyde 3-phosphate dehydrogenase.

### Microscope imaging

Wide-field images were acquired on an Upright Leica microscope (model DMR) fitted with an Orca-100 digital camera (model C4742-95; Hamamatsu Photonics) and a 40x 1.0 numerical aperture (NA) oil Plan Fluotar objective. Apotome images were acquired using an epifluorescence microscope system (AxioVision Z1; Carl Zeiss, Thornwood, NY) fitted with an Apotome slide module, AxioCam MRm digital camera, and a 40x 0.5 EC Plan-Neofluar or 100x 1.4 NA oil Plan-Apochromat objective (Carl Zeiss). For each experiment, images were acquired using the same imaging conditions.

Confocal z-stacks (z-step size of 0.5 µm) of whole mount samples were acquired using a Nikon A1R confocal laser microscope equipped with GaAsP detectors and a 60× Plan-Apochromat objective lambda with a NA of 1.4 and run by NIS Elements software (Nikon). NIS Elements (version 5.02) was used to generate 3D reconstructions of z-stacks using the Volume Viewer tool with z-depth coding blending and the Rainbow contrast look up table and to determine the z position of the melanocyte cell body centroid.

### Fluorescence intensity of cell-cell borders

For each image, background intensity was defined as the average of three mean pixel intensity measurements from an image without fluorescence. Fluorescence pixel intensity at random KC borders surrounding a MC or a cluster of MCs was determined by measuring the mean pixel intensity at a defined border and dividing by the border length.

### Quantification of dendrite length

MCs were grown to 75% confluence in MC media and subsequently switched to a 1:1 ratio of KC conditioned media or continued to be grown in MC media alone. Cells were incubated overnight (12hrs) or for 7 days before being fixed in 4% paraformaldehyde (Sigma-Aldrich) in PBS for 10 minutes. Wide-field images were used to visualize MC dendrites following incubation. Dendrite parameters were quantified with Fiji software (NIH). Dendrites were defined as extensions originating from the cell body that did not exhibit a lamellar morphology. Dendrite length was measured from cell body to the tip of the dendrite (in µm). When applicable, Reparixin (Moriconi *et al.*, 2007) (Cayman Chemical, Ann Arbor, MI) was added to media at a final concentration of 5 µg/ml media. Recombinant IL6 and IL8 (Abcam) were added to media at final concentrations of 0.2 ng/ml and 25 ng/ml, respectively.

### Quantification of melanin secretion

MCs were grown to confluence in MC media and subsequently switched to a 1:1 ratio of KC conditioned media or media alone with fresh media added to existing media every 2 days up to 7 days. After 7 days, media was collected and subjected to centrifugation at 11,000 rpm for 1 minute. The resulting melanosome pellet was re-suspended in 200 µl PBS and transferred to the appropriate well of a 96-well plate (BD Biosciences). The absorbance at 405 nm of each well was measured with an ELx800 microplate reader (Bio-Tek Instruments, Inc). Absorbance readings were plotted after normalizing to a media only control for each condition in 3 independent experiments. When applicable, 100 nM Recombinant Human Beta Defensin 3 (Sigma Aldrich, Saint Louis, MO) was used to inhibit MC1R (Swope *et al.*, 2012; Wolf Horrell *et al.*, 2016).

### Sample size and statistical analysis

All experiments were performed independently ≥3 times (i.e. biological replicates performed on different days, not technical replicates performed in parallel at the same time). For each independent/biological replicate, multiple experimental/control arms were processed and analyzed in parallel. Representative experiments are displayed throughout the figures. All graphs are displayed as mean ± standard error of the mean (SEM). Power analyses to determine sample sizes were determined in consultation with the Northwestern University Quantitative Data Sciences Core. Two group comparisons were performed using two-tailed, two-sample equal variance Student’s t test. For comparisons of more than two groups, One-way ANOVA on ranks (Kruskal-Wallis) was used, followed by Tukey or Dunn’s multiple comparisons test. All analysis was performed using GraphPad Prism version 7.0 for Mac (GraphPad Software, La Jolla, CA, USA). Statistical significance was defined as, *p<0.03, **p<0.002, ***p<0.0002, ****p<0.0001.

## Acknowledgements

This work was supported by NIH/NIAMS R01 AR041836 and NIH/NCI R01 CA122151 to KJG and by the Liz and Eric Lefkofsky Family Foundation Innovation Research Award to KJG and JLJ. Additional support was provided by the JL Mayberry endowment to KJG. CRA was supported through a NIH/NCI Ruth L. Kirschstein Training Grant through Northwestern University’s Robert H. Lurie Comprehensive Cancer Center (T32 CA070085-14) “Signal Transduction in Cancer” and also through a NIH/NCI Ruth L. Kirschstein National Research Service Award 1F32CA210498-01. We thank Dr. Zalfa Abdel-Malek (University of Cincinnati) for critical reading of the manuscript. We acknowledge support and materials from the Northwestern University Skin Disease Research Center supported by 5P30AR057216. Imaging work was performed at the Northwestern University Center for Advanced Microscopy generously supported by NCI CCSG P30 CA060553 awarded to the Robert H Lurie Comprehensive Cancer Center.

## Abbreviations

KC: Keratinocyte
MC: Melanocyte
Dsg1: Desmoglein 1
UV: Ultraviolet light
E-cad: Epithelial cadherin
SAM: Severe dermatitis, multiple allergies, and metabolic wasting

